# Histopathological Effects of Therapeutic Doses of Combined XO-Inhibitors and ACE-Inhibitors on the Expression of VEGF-A in the Myocardium and Renal Cortex in Chronic Hypertensive Albino Rats

**DOI:** 10.1101/163824

**Authors:** Hassan M. Rezk, Afaf Ibraheim

## Abstract

**Background:** Hypertension is risk factor for development of congestive heart failure. The pathogenesis of myocardial and renal cortex changes in hypertension includes structural remodeling and fibrosis.

**Aim of study:** is to evaluate the effects of therapeutic doses of combined XO-Inhibitors and ACE-Inhibitors on expression of VEGF-A in the myocardium and renal cortex in chronic hypertensive albino rats.

**Material & Methods:** Thirty male albino rats were divided into: Group I: (control group), Group II (Non-treated Hypertensive rats), Group III (Allopurinol-treated Hypertensive rats), Group IV (Captopril-treated Hypertensive rats) and Group V (Allopurinol-Captopril-treated Hypertensive rats). At 4 and 8 weeks, the rats were anesthetized followed by obtaining of heart and left kidney to be stained with Masson trichrome and Anti-Vascular endothelial growth factor-A antibody.

**Results:** Group II, one month hypertensive rats showed, myocardium showed disarray with significant increase in interstitial fibrosis. The renal cortex showed evidences indicating glomerulosclerosis. Immunohistochemistry, there was high significant decrease in the number of cells of renal cortex with +ve expression of VEGF-A. Later, they showed marked significant increase in interstitial fibrosis. In Group III, myocardium and renal cortex showed high significant increase in interstitial fibrosis. After two month, there were high significant decrease in the surface area of interstitial fibrosis in myocardium and renal cortex with high significant decrease number of the myocardium with +ve expression of VEGaF-A. In Group IV, myocardium showed disarray with marked significant reduction in interstitial fibrosis. The renal cortex showed marked significant reduction in the interstitial fibrosis with significant decrease in the number of cells with +ve expression of VEGF-A. Later, myocardium showed the most high marked significant reduction in interstitial fibrosis with highly significant increase in number of cells with positive expression of VEGF-A. In Group V after two month, both myocardium and renal cortex showed nearly normal architecture with marked significant reduction in interstitial fibrosis.

**Conclusions:** Long term therapy with the combination between allopurinol and captopril decreases the fibrotic changes associated with hypertension and enhances the process of angiogenesis.

## 1. Introduction

Reactive oxygen species (ROS) (e.g. superoxide anion (O2**^-^**)) are known to be involved in many diseases and disorders including hypertension and ischemia which are known as ROS related diseases. Serum lipid peroxides or ROS are increased in essential hypertensive patients or hypertensive animal models.^1^

Several studies have focused on XO as a source of oxidative stress. Xanthine oxidase (XO) plays important role in excessive generation of ROS especially superoxide anion (O2**^-^**). XO is known to exist initially as xanthine dehydrogenase in many tissues, it is converted to XO. Its activity being highly upregulated in various pathological conditions, which becomes a major source of (O2**^-^**). As a consequence, (O2**^-^**) generation by XO is regarded as an etiological mechanism of such diseases, and thus it is considered as a therapeutic target for these ROS related diseases^2^.

Xanthine oxidoreductase, when transferred into its oxidase form xanthine oxidase (XO), generates super-oxide and hydrogen peroxide upon conversion of xanthine to uric acid. Vascular and myocyte-localized XO is increased in coronary artery disease, and circulating XO levels are upregulated in congestive heart failure. ^3^

Growing evidence supports the role of uric acid as a biomarker of oxidative stress and as a mediator of hypertension.^4^ Several evidences have been shown positive relationship between serum uric acid (UA) levels and cardiovascular mortality in patients with chronic kidney disease. The reduction in serum UA levels by administering the xanthine oxidase (XO) inhibitor allopurinol has been shown to slow the progression of renal dysfunction and decrease the risk of cardiovascular disease. This beneficial effect is thought to originate from a lowering of the plasma UA level, because UA itself has been shown to generate oxidative stress in adipocytes, vascular endothelial cells, and vascular smooth muscle cells.

XO inhibition in hyperuricemic with dilated and ischemic cardiomyopathy leads to an improvement of vascular Nitric Oxide (NO) bioavailability^5^, and local infusion of allopurinol into the coronary circulation in patients with dilated cardiomyopathy lowered myocardial oxygen consumption.^6^

Allopurinol is a potent xanthine oxidase inhibitor that is used in hyperuricemic patients to prevent gout. It has also been shown to decrease cardiovascular complications and to reduce blood pressure in hypertensive patients.^7^

Harrison et al. ^8^have found that angiotensin II (AII) administration markedly increases ROS. This effect is suppressed by AII receptor antagonists. These findings strongly suggest that ROS are involved in AII-induced blood pressure elevation or vascular damage.^9^ Angiotensin II-induced endothelial dysfunction also shows the significant role of XO in oxidative injury.

Endothelium preserves its integrity through endothelium-relaxing dependent factor, which is the best to be characterized as NO. ^10^ Therefore, NO plays an important role in the regulation of blood pressure^11^ Impaired NO bioavailability will result in reduced endothelium-dependent vasorelaxation, eventually leading to hypertension.^12^

Vascular oxidative stress increased production of ROS with vascular dysfunction. It leads to imbalance between the activity of endogenous pro-oxidative enzymes e.g. xanthine oxidase and anti-oxidative enzymes e.g. superoxide dismutase in favor of the former. Increased ROS concentrations reduce the amount of bioactive Endothelium-derived (NO) by chemical inactivation to form toxic per-oxy-nitrite. Enhanced inactivation and/or reduced synthesis of NO is seen in conjunction with risk factors for cardiovascular disease. Therapeutically, drugs in clinical use such as ACE inhibitors can improve endothelial function.^13^

Angiotensin-converting enzyme (ACE) plays a vital role in the regulation of BP via the active form, angiotensin II (Ang II). ACE is mainly located on the surface of the endothelium and epithelium involved in the potent vasoconstriction of blood vessels, subsequently leading to elevation of BP. Ang II also stimulates the release of aldosterone, further increasing blood volume and BP due to water and salt retention. ^14^

Angiotensin-converting enzyme inhibitors (ACE-I) were developed as therapeutic agents targeted for the treatment of hypertension. Captopril is the prototype of the sulfhydryl-containing ACE inhibitors^[15]^. Sulfhydryl angiotensin-converting enzyme inhibitors induce sustained reduction of systemic oxidative stress and improve the nitric oxide pathway in patients with essential hypertension.^16^

The effects of antihypertensives like angiotensin converting enzyme (ACE) inhibitor and Ca channel blocker are in part due to antioxidant activity. ACE inhibitors increase NO availability by reducing angiotensin II production and bradykinin degradation. Antihypertensive therapy taking this perspective into account would most likely be effective to prevent complications of hypertension.^17^

In 1934, Goldblatt et al. developed a hypertension model through partial constriction of the renal artery in dog. This has led to induction of hypertension model using rats, rabbits, sheep, and cats.^18^ When the renal artery is ligated or constricted, Renin–Angiotensin–Aldosterone System (RAAS) is activated. Angiotensinogen is converted to angiotensin-I (Ang I) in the presence of renin secreted by kidney.

ROS regulate collagen metabolism in a variety of non-cardiac cell types. However, it is not known whether ROS can regulate collagen metabolism in cardiac fibroblasts, which are responsible for collagen synthesis and degradation in the myocardium ^19^. Renal interstitial fibrosis is one of the common histopathological features of progressive renal disease with diverse etiology.^20^

Vascular endothelial growth factor (VEGF) is a glycoprotein expressed in multiple organs which plays a key role in maintaining homeostasis and cell survival. The gene undergoes alternative splicing, and six VEGF isoforms have been identified, with the most biologically active variant being VEGF165 (VEGF-A).^21^ VEGF is the most potent and primary endothelial specific angiogenic growth factor, both in physiological and pathological conditions through VEGF signaling pathway. ^22^ This pathway often seems to be affected by ROS. VEGF induces endothelial cell migration and proliferation through an increase of intracellular ROS. ^23^ Specific cells that express VEGF include progenitor endothelial cells, endothelial cells (EC), podocytes (renal epithelial cells), fibroblasts, macrophages.

Vascular endothelial growth factor A (VEGF-A) is the dominant inducer to the growth of blood vessels. VEGF-A is essential for adults during organ remodeling and diseases that involve blood vessels. VEGF-A increase the endothelial permeability and swelling and also stimulating angiogenesis. Once released, VEGF-A cause the cell to survive, move, or further differentiate.^24^

The aim of this study is to evaluate the effects of therapeutic doses of combined XO-Inhibitors and ACE-Inhibitors on expression of VEGF-A in the myocardium and renal cortex in chronic hypertensive albino rats.

## 2. Materials and Methods

### 2.1 Animals

Thirty male albino rats (240 - 280 g) were used in the present study. The animals were housed in cages at room temperature (22–25°C) and in a photoperiod of 14-h light/10-h dark/day. Rats were maintained on standard laboratory balanced commercial diet and water ad libitum. Rats were obtained from and housed in the animal house, faculty of pharmacy, Mansoura University, Egypt. All experiments were performed in line with the ethical considerations, recommended by the Faculty of Medicine, Mansoura University, Egypt.

### 2.2. Experimental design

The animals were divided into five groups

#### a. Group I

Six rats were served as a control group receiving 0.5 mL of saline by injection.

#### b. Group II (Non-treated Hypertensive rats)

Six rats with induced hypertension through clipping of right renal artery did not receive any medications throughout the period of the experiments (8 weeks).

#### c. Group III (Allopurinol-treated Hypertensive rats)

Six rats with induced hypertension through clipping of right renal artery and treated with therapeutic dose of Allopurinol started when rats’ systolic blood pressure reached 150 mmHg 3 weeks postoperative for 8weeks.

#### d. Group IV (Captopril-treated Hypertensive rats)

Six rats with induced hypertension through clipping of right renal artery and treated with therapeutic dose of Captopril tablets started when rats’ systolic blood pressure reached 150 mmHg 3 weeks postoperative for 8weeks.

#### e. Group V (Allopurinol-Captopril-treated Hypertensive rats)

Six rats with induced hypertension through clipping of right renal artery and treated with therapeutic doses of allopurinol and Captopril started when rats’ systolic blood pressure reached 150 mmHg 3 weeks postoperative for 8weeks.

### 2.3. Protocol for right renal artery clipping

The experimental rats were underwent an operation of renal artery constriction. Under anesthesia with 3% sodium pentobarbital (36 mg/kg intraperitoneal), a median longitudinal incision on abdominal skin was performed, then a ring-shaped silver clip with an inner diameter of 0.30 mm was placed around the root of right renal artery, but the left contralateral kidney remained untouched. All postoperative care were done for all experimental rats. All rats were allowed an ordinary rat chow diet (plant protein 15.9%, nonfish animal protein 5.4%, fish protein 1.7%; carbohydrate 52.5%; unsaturated fat 3.4%, saturated fat 1.3%; Na1 0.24%, K1 1.0%) and tap water as desired and kept on a 12-hour light/dark cycle.^25^

### 2.4. Protocol for Allopurinol and Captopril intake

Allopurinol (Zyloric, 300 mg, GlaxoSmithKline, gsk) and Captopril tablets (Capoten, 25 mg, SmithKline Beecham, Egypt, L.L.C.) were dissolved in drinking water ad libitum. The dose of allopurinol per day was carefully calculated by the daily water intake and body weight was approximately 3 mg/kg/day to avoid possible renal damages due to xanthine calculi formation. The dose of captopril was (60 mg/kg/day)^26^ started when rats’ systolic blood pressure reached 150 mmHg 3 weeks postoperative for 8weeks.The allopurinol and captopril treatment were started as soon as the rats were chronically hypertensive.^27^

### 2.5. Measurement of the blood pressure in rats

Systolic blood pressure (SBP) was measured using a tail-cuff sphygmomanometer in pharmacology department, faculty of pharmacy, Mansoura University, Egypt. All animals were acclimated for blood pressure measurements 1 week before and weekly intervals after renal artery clip for during the drug treatment for 8 weeks. The mean systolic BP in rats was 110 mm Hg before renal artery constriction. The experimental rats were started to receive allopurinol and captopril when the hypertension becomes chronic, i.e. blood pressure became 150 mm Hg (after 3 postoperative weeks). The experimental rats did not show neither renal failure nor renal failure during our study. ^28^

### 2.6. Histopathological Immunohistochemical Studies

At the assigned time of scarification 4 and 8 weeks, the rats were anesthetized by pentobarbital overdose (200mg/kg) inhalation followed by mid-sternal incision followed by obtaining of heart and left kidney where they were placed in 10% formaldehyde. All the specimen were removed and prepared for paraffin blocks in pathology department, faculty of Medicine, Mansoura University, Egypt. Sections were cut with a microtome (Leica RM 2025, Germany) at 5 μm thicknesses and stained with Hematoxylin and Eosin as a routine histological technique and Masson trichrome for staining of collagen fibers indicator for the degree of fibrosis.

### 2.7. Immunohistochemical Study

Vascular endothelial growth factor-A (VEGF-A) in the myocardium and renal cortex were detected by immunohistochemistry. Sections were incubated with the polyclonal primary antibody against VEGF-A (Anti-VEGF-A antibody) (Sigma is now MERCK, AB1876-I EMD MILLIPORE). All sections were counterstained with Eosin, dehydrated, mounted, and viewed by light microscopy. The brown color stained cells were considered positive expression of VEGF-A.

### 2.8. Computer Assisted digital image analysis (Digital morphometric study)

Slides stained with Masson trichrome and Anti-VEGF-A antibodies were photographed using Olympus® digital camera installed on Olympus® microscope with 1/2 X photo adaptor, using 40 X objective. Four sections from each stain for heart and kidney were used. The result images were analyzed on Intel® Core I3® based computer using Video-Test Morphology® software (Russia) with a specific built-in routine for measurement of the collagen surface areas (Masson trichrome stained areas) and the number of positive anti-VEGF-A antibody cells in myocardium and renal cortex after 4 and 8 weeks according to the planned groups.

### 2.9. Statistical Analysis

Data from Masson trichrome stained and anti-VEGF-A antibody stained sections from all groups were analyzed using Statistical Package for Social Science software computer program version 17 (SPSS, Inc., Chicago, IL, USA). Quantitative parametric data were presented in mean and standard deviation, while Quantitative non- parametric data (the data with abnormal distribution of Anti-VEGF-A antibody rats’ hearts of one month hypertensive with or without treatment stained sections) were presented in median and interqurtile range (IQR). For quantitative parametric data, one way Analysis of variance (ANOVA) and tukey were used for comparison of different groups and student’s t-test (Paired) for comparing two related groups while for comparing quantitative non-parametric data, Kruskal Wallis test followed by Mann Whitney comparison were used for comparison of different groups and Wilcoxon signed rank test was used for comparing two related groups. P value less than 0.05 was considered statistically significant.

## 3. Results

### 3.1. After 4 weeks

The myocardium in Control Group I showed normal regular striations with significant absence of interstitial fibrosis. The renal cortex in the same group, appeared normal with normal glomeruli. Some areas of renal cortex showed significant fibrotic areas (Figs 1, 15 & 16) and (Tables 1 & 2). In immunohistochemical study of group I, some cardiac muscle fibers showed positive expression of VEGF-A. There was high significant number of cells of glomeruli and renal tubules with positive expression of VEGF-A. (Figs 10, 17 & 18) and (Tables 3 & 4).

**Table 1.**
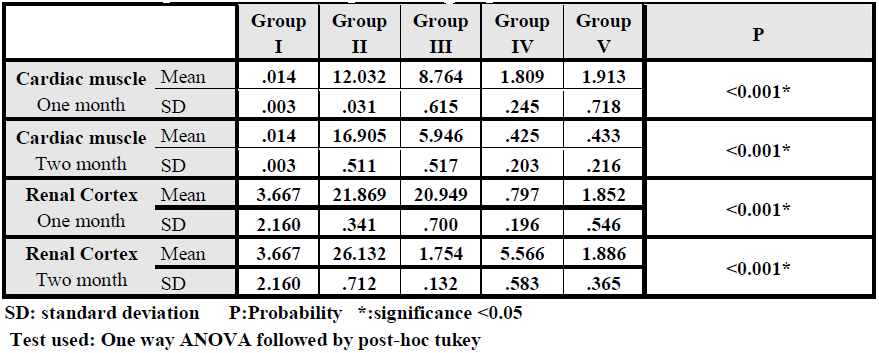
Comparison between collagen surface area (Masson Trichrom stain) in cardiac muscle and renal cortex among studied control & experimental groups in one and two months.

**Table 2.** Comparison between collagen surface area (Masson Trichrom stain) in one & two months in cardiac muscle and renal cortex in studied experimental groups

|  |  | One month |  | Two month |  | P |
| --- | --- | --- | --- | --- | --- | --- |
|  |  | Mean | ±SD | Mean | ±SD |  |
| Cardiac muscle | Group II | 12.032 | .031 | 16.905 | .511 | <0.001* |
|  | Group III | 8.764 | .615 | 5.946 | .517 | .001* |
|  | Group IV | 1.809 | .245 | .425 | .203 | <0.001* |
|  | Group V | 1.913 | .718 | .433 | .216 | .01* |
| Renal Cortex | Group II | 21.869 | .341 | 26.132 | .712 | <0.001* |
|  | Group III | 20.949 | .700 | 1.754 | .132 | <.001* |
|  | Group IV | .797 | .196 | 5.566 | .583 | <0.001* |
|  | Group V | 1.852 | .546 | 1.886 | .365 | 0.9 |

SD: standard deviation P:Probability \*:significance <0.05
Test used: Student's t-test(Paired)

**Table 3.**
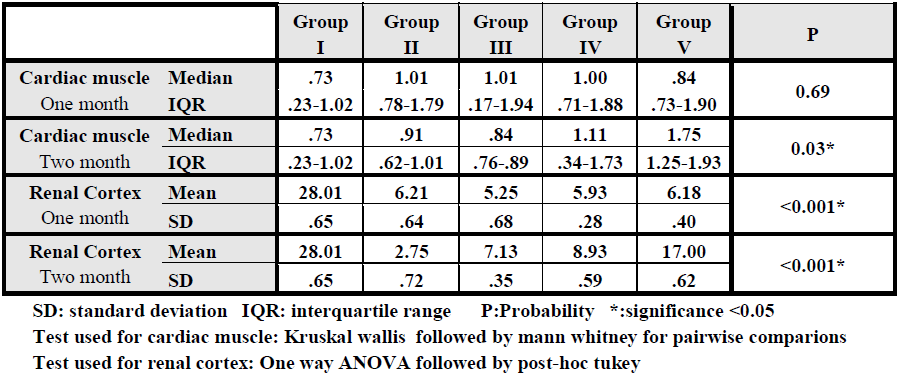
Comparison between the number of positive anti-VEGF-A antibody in cardiac muscle and renal cortex among studied control & experimental groups in one and two months.

**Table 4.** Comparison between the number of positive anti–VEGF A antibody in one & two months in cardiac muscle and renal cortex in studied experimental groups

|  |  | One month |  | Two month |  | P |
| --- | --- | --- | --- | --- | --- | --- |
|  |  | Median | IQR | Median | IQR |  |
| Cardiac muscle | Group II | 1.01 | .78-1.79 | .91 | .62-1.01 | 0.46 |
|  | Group III | 1.01 | .17-1.94 | .84 | .76-.89 | 0.34 |
|  | Group IV | 1.00 | .71-1.88 | 1.11 | .34-1.73 | 0.9 |
|  | Group V | .84 | .73-1.90 | 1.75 | 1.25-1.93 | 0.075 |
|  |  | Mean | ±SD | Mean | ±SD |  |
| Renal Cortex | Group II | 6.21 | .64 | 2.75 | .72 | <0.001* |
|  | Group III | 5.25 | .68 | 7.13 | .35 | .003* |
|  | Group IV | 5.93 | .28 | 8.93 | .59 | <0.001* |
|  | Group V | 6.18 | .40 | 17.00 | .62 | <0.001* |

SD: standard deviation IQR: interquartile range P:Probability \*:significance <0.05
Test used for cardiac muscle: Wilcoxon signed rank test
Test used for renal cortex: Student's t-test(Paired)

**Figure 1:**
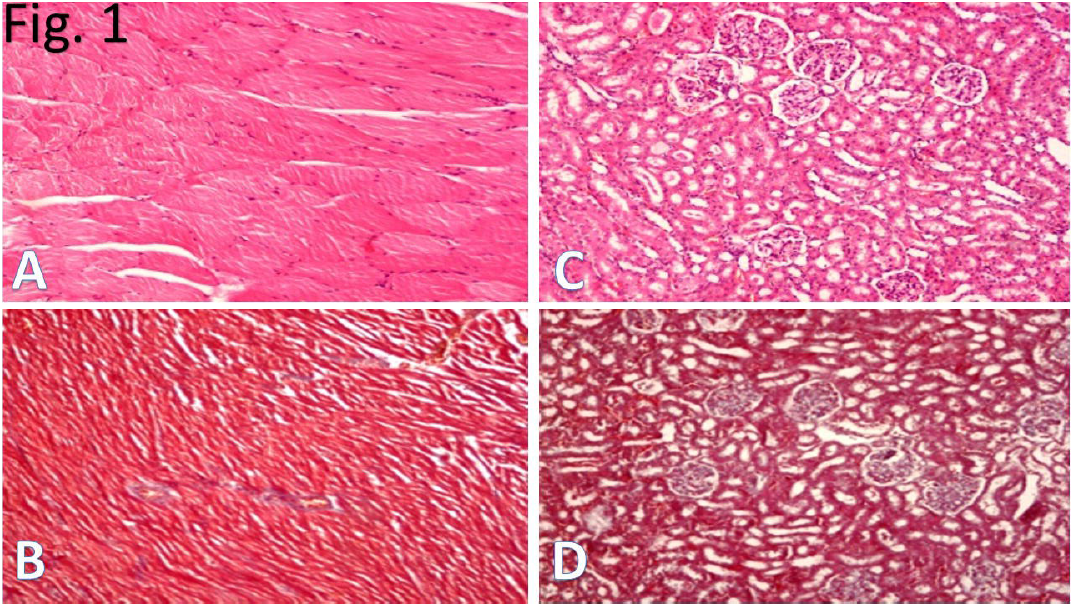
**(A, B, C & D):** Photomicrographs of histological sections of heart (A & B) and kidney (C & D) of control group (Group I) in one and two month. **A & B:** cardiac muscle showed normal architecture with regular striations. **C & D:** renal cortex showed normal structure with normal glomeruli. Some areas of renal cortex showed fibrotic areas. **(A & C: Hx. & E. stained sections (X 100), B & D: Masson Trichrome stained sections (X 100))**

**Figure 10:**
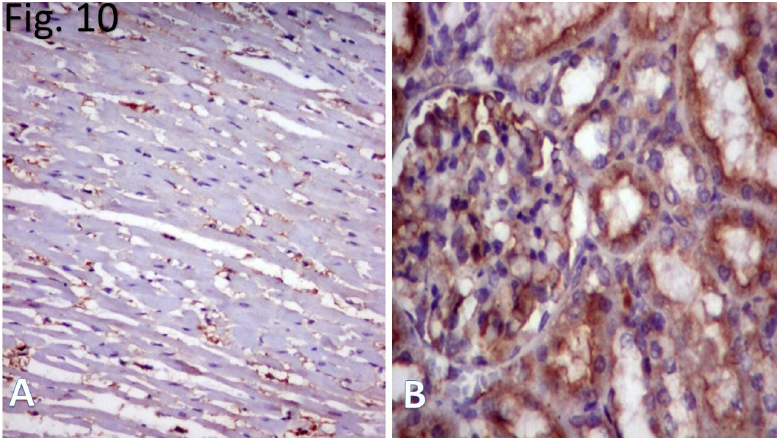
**(A & B):** Photomicrographs of histological sections of heart (A) and kidney (B) of control group (Group I) in one and two months. **A:** some cardiac muscles showed positive expression of VEGF-A. **B:** most of cells of glomeruli and renal tubules showed positive expression of VEGF-A. **(Immunohistochemical stain, Counterstained with E., X400)**

**Figure 15:**
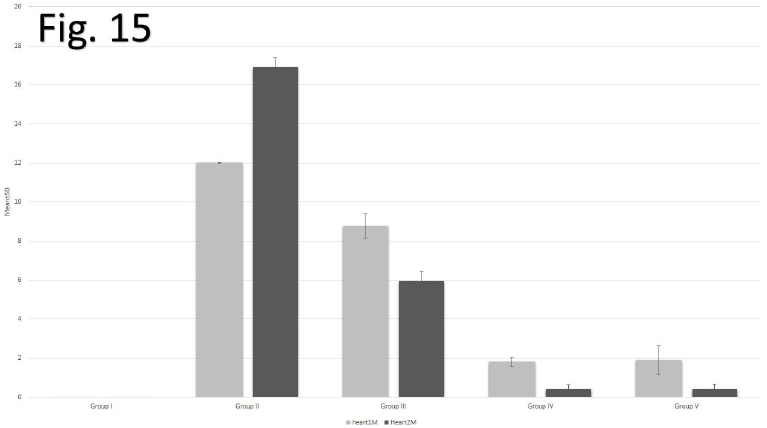
A chart showed the comparison between collagen surface area (Masson Trichrom stain) in cardiac muscle among studied control & experimental groups in one and two months.

**Figure 17:**
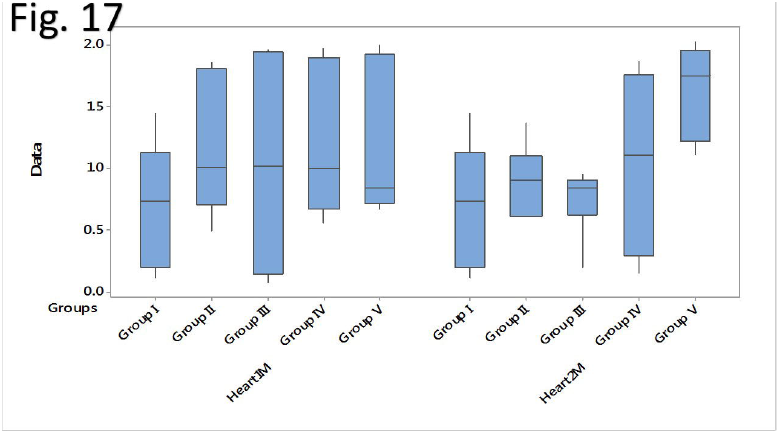
A chart showed the comparison between the number of positive anti-VEGF-A antibody in cardiac muscle among studied control & experimental groups in one and two months.

**Figure 18:**
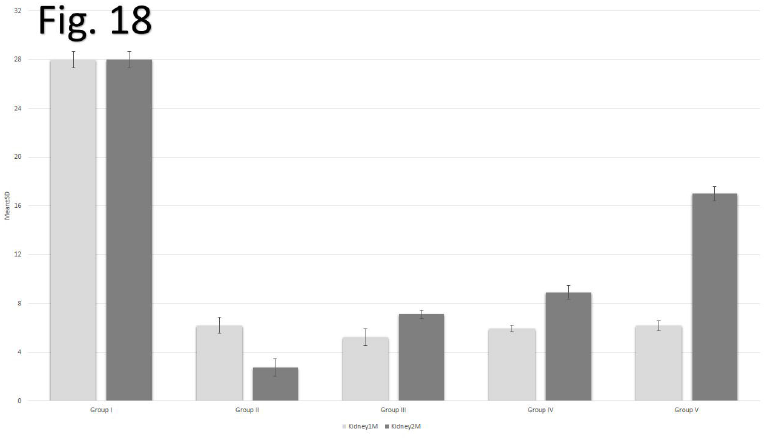
A chart showed the comparison between the number of positive anti-VEGF-A antibody in renal cortex among studied control & experimental groups in one and two months.

In histopathological examination of Group II (Non-treated Hypertensive rats), myocardium showed disarray with significant increase in interstitial fibrosis. The renal cortex showed ischemic degenerative changes in glomeruli and renal tubules with significant increase in interstitial fibrosis indicated glomerulosclerosis. (Figs 2, 15 & 16) and (Tables 1 & 2). The anti-VEGF-A antibody stained sections, non-significant increase in the number of cardiac muscle fibers with positive expression of VEGF-A in comparison with that of the control group. While, there was high significant decrease in the number of cells of renal cortex with positive expression of VEGF-A. in comparison with that of the control group (Figs 11, 17 & 18) and (Tables 3 & 4).

**Figure 2:**
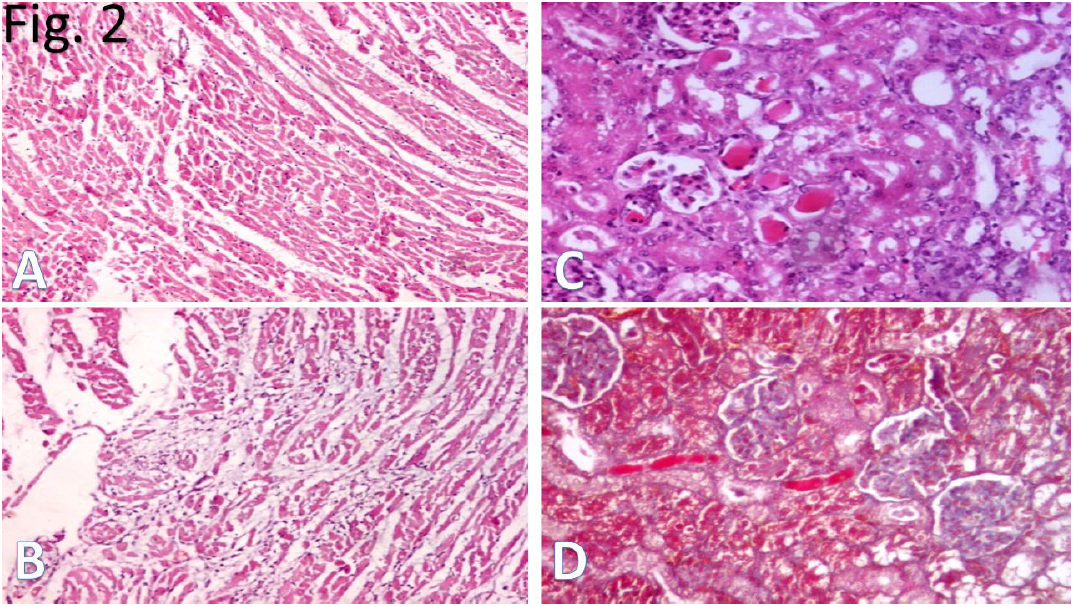
**(A, B, C & D):** Photomicrographs of histological sections of heart (A & B) and kidney (C & D) of Non-treated Hypertensive rats (Group II) for one month. **A:** cardiac muscle showed disarray of cardiac muscle fibers **(Hx. & E., X100). B:** cardiac muscle showed increased interstitial fibrosis **(Masson trichrome, x100)**. **C:** renal cortex showed ischemic degenerative changes in glomeruli and renal tubules **(Hx. & E., X100)**. **D:** renal cortex showed increased interstitial fibrosis **(Masson trichrome, x100).**

**Figure 11:**
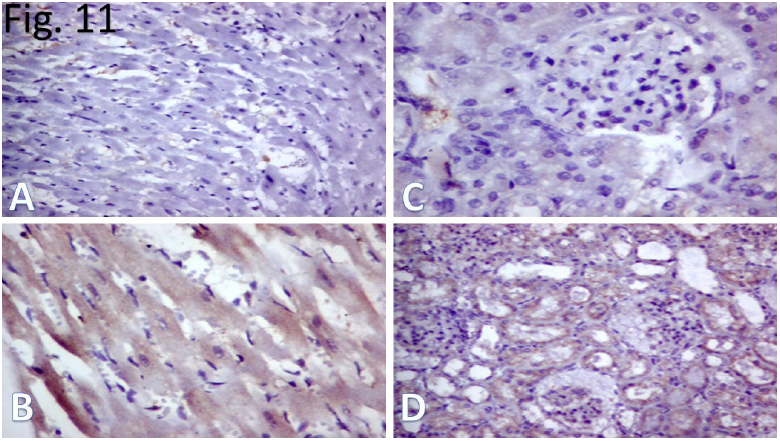
**(A, B, C & D):** Photomicrographs of histological sections of heart (A & B) and kidney (C & D) of Non-treated Hypertensive rats (Group II). **After one month, A:** some cardiac muscle fibers showed positive expression of VEGF-A. **C:** some cells of renal cortex showed positive expression of VEGF-A. **After two month, B:** few cardiac muscle showed positive expression of VEGF-A. **D:** few cells of renal cortex renal tissue showed positive expression of VEGF-A. **(Immunohistochemical stain, Counterstained with E., X400)**

In Group III (Allopurinol-treated Hypertensive rats), the histopathology of myocardium showed disarray with high significant increase in interstitial fibrosis in comparison to that of the control otherwise it is less than that of group II for the same duration. The renal cortex showed mild ischemic degenerative changes in glomeruli and renal tubules. It showed high significant increase in interstitial fibrosis the same as that of the group II for the same duration (Figs 4, 15 & 16) and (Tables 1 & 2). Non-significant number of the cardiac muscle fibers showed positive expression of VEGF-A in comparison to groups I & II. In group III, renal cortex still had shown significant decrease in the number of cells with positive expression of VEGF-A in comparison to group I but it was similar to that of group II. (Figs 12, 17 & 18) and (Tables 3 & 4).

**Figure 4:**
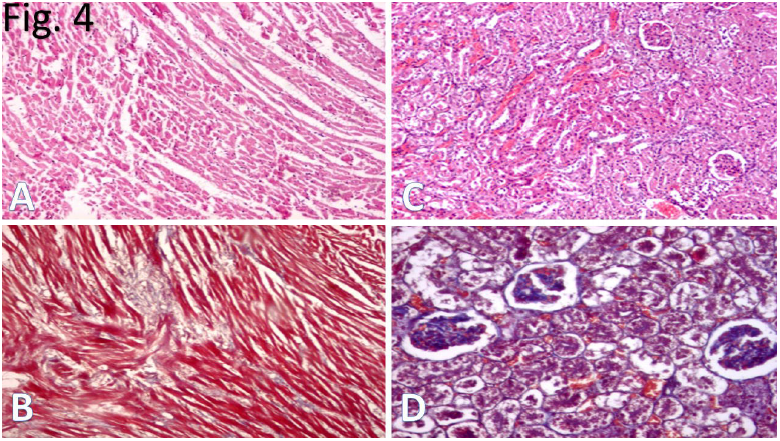
**(A, B, C & D)** Photomicrographs of histological sections of heart (A & B) and kidney (C & D) of Allopurinol-treated Hypertensive rats (Group III) for one month. **A:** cardiac muscle showed with disaary of cardiac muscle fibers **(Hx. & E., X100). B:** cardiac muscle showed increased in interstitial fibrosis **(Masson trichrome, X200)**. **C:** renal cortex showed mild ischemic degenerative changes in glomeruli and renal tubules **(Hx. & E., X100) D:** renal cortex showed marked increase in interstitial fibrosis and thickened basement membrane **(Masson trichrome, X200)**.

**Figure 12:**
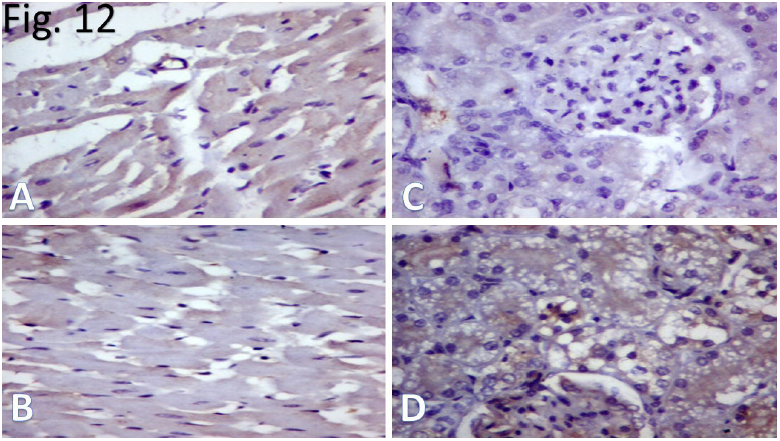
**(A, B, C & D)** Photomicrographs of histological sections of heart (A & B) and kidney (C & D) of Allopurinol-treated Hypertensive rats (Group III). **After one month, A:** most of cardiac muscle showed positive expression of VEGF-A. **C:** some cells of renal cortex showed positive expression of VEGF-A. **After two month, B:** few of cardiac muscle showed positive expression of VEGF-A. **D:** some cells of renal cortex renal tissue showed positive expression of VEGF-A. **(Immunohistochemical stain, Counterstained with E., X400)**

In Captopril-treated Hypertensive rats (Group IV), myocardium still showed disarray with marked significant reduction in interstitial fibrosis in comparison to groups II & III of the same duration. The renal cortex showed atrophy of some glomeruli with marked significant reduction in the interstitial fibrosis in comparison to groups I, II & II of the same duration (Figs 6, 15 & 16) and (Tables 1 & 2). As in groups I, II & III, the number cardiac muscle fibers with positive expression of VEGF-A was still non-significant. The renal cortex showed significant decrease in the number of cells with positive expression of VEGF-A. in comparison to group I with slight increase than that of group III. (Figs 13, 17 & 18) and (Tables 3 & 4).

**Figure 6:**
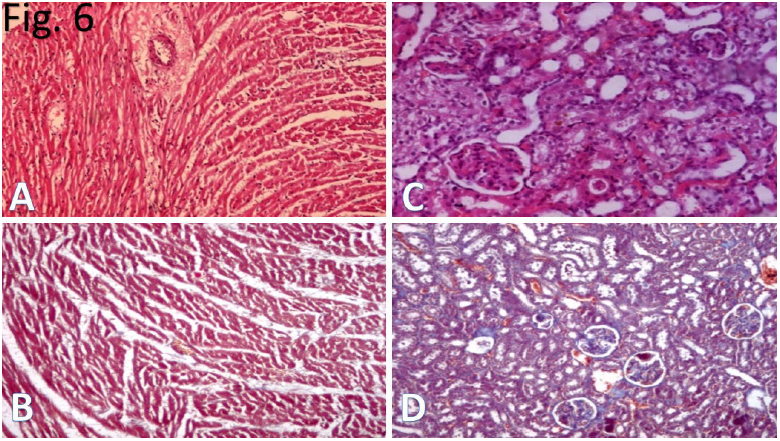
**(A, B, C & D)** Photomicrographs of histological sections of heart (A & B) and kidney (C & D) of Captopril-treated Hypertensive rats (Group IV) for one month. **A:** cardiac muscle still showed disarray of cardiac muscle fibers **(Hx. & E., X100). B:** with mild interstitial fibrosis **(Masson trichrome, X200). C:** renal cortex showed atrophy of some glomeruli **(Hx. & E., X100) D:** renal cortex showed mild interstitial fibrosis and thickened basement membrane **(Masson trichrome, X200).**

**Figure 13:**
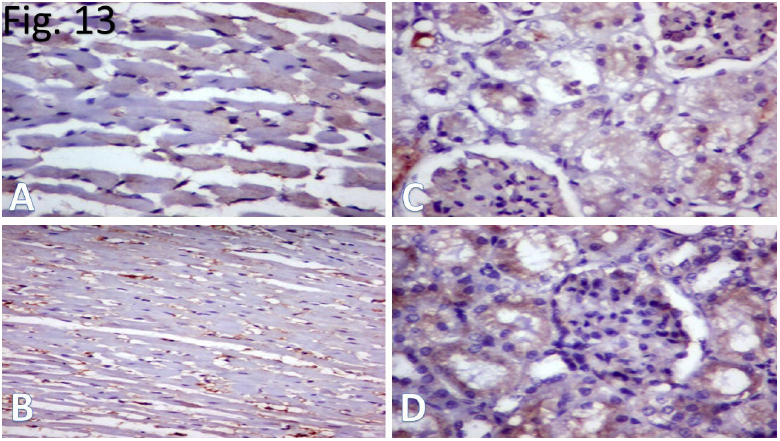
**(A, B, C & D)** Photomicrographs of histological sections of heart (A & B) and kidney (C & D) of Captopril-treated Hypertensive rats (Group IV). **After one month, A:** some of cardiac muscle showed positive expression of VEGF-A. **C:** some cells of renal cortex showed positive expression of VEGF-A. **After two month, B:** most of cardiac muscle showed positive expression of VEGF-A. **D:** some cells of renal cortex renal tissue showed positive expression of VEGF-A. **(Immunohistochemical stain, Counterstained with E., X400)**

In Group V (Allopurinol-Captopril-treated Hypertensive rats), the histopathological examination of myocardium still showed mild disarray with significant reduction in interstitial fibrosis with the same that of the of group IV in comparison to groups II & III. The renal cortex showed normal structure of renal cortex and with normal glomeruli with marked reduction in fibrosis the same as in group IV in comparison to groups II & III (Figs 8, 15 & 16) and (Tables 1 & 2). The immunohistochemical stained sections, there was non-significant increase in the number of cardiac muscle fibers with positive expression of VEGF-A.in comparison to groups I, II, III & IV for the same duration. The renal cortex still had showed significant decrease in the number of cells with positive expression of VEGF-A. in comparison to group I with slight increase than that of group III & IV. (Figs 14, 17 & 18) and (Tables 3 & 4).

**Figure 8:**
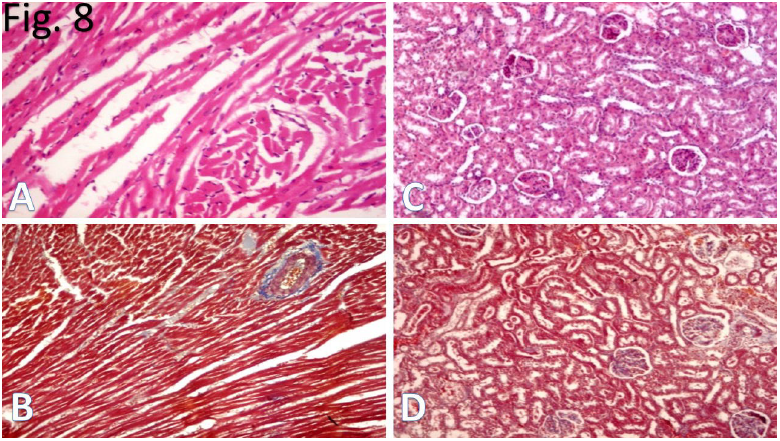
**(A, B, C & D)** Photomicrographs of histological sections of heart (A & B) and kidney (C & D) of Allopurinol-Captopril-treated Hypertensive rats (Group V) for one month. **A:** cardiac muscle still showed mild disarray of cardiac muscle fibers **(Hx. & E., X100). B:** cardiac muscle showed mild increase in interstitial fibrosis **(Masson trichrome, X200). C & D** showed normal structure of renal cortex and with normal glomeruli with minimal fibrosis. **(Hx. & E., X200 & Masson trichrome, X200).**

**Figure 14:**
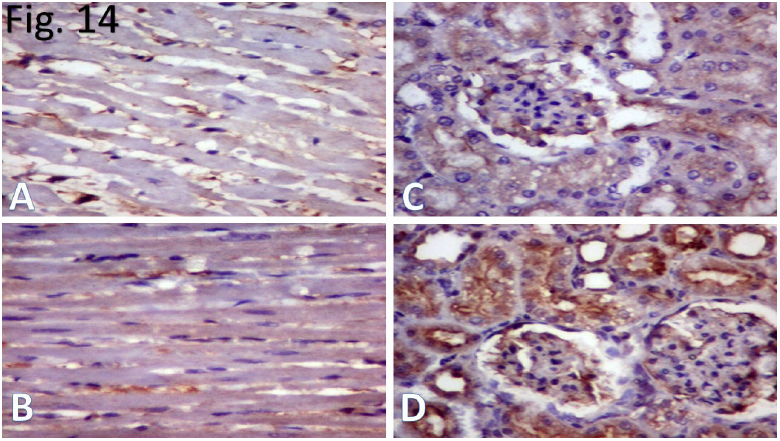
**(A, B, C & D)** Photomicrographs of histological sections of heart (A & B) and kidney (C & D) of Allopurinol-Captopril-treated Hypertensive rats (Group V). **After one month, A:** some of cardiac muscle showed positive expression of VEGF-A. **C:** some cells of renal cortex renal tissue showed positive expression of VEGF-A. **After two month, B:** some of cardiac muscle showed positive expression of VEGF-A. **D:** most cells of renal cortex renal tissue showed positive expression of VEGF-A. **(Immunohistochemical stain, Counterstained with E., X400)**

### 3.2. After 8 weeks

The myocardium and renal cortex in Control Group I for two month showed the same histopathological and immunohistochemical findings as the same group for one month normal architecture with regular striations with significant absence of interstitial fibrosis (Figs 1, 10, 15, 16, 17 & 18) and (Tables 1, 2, 3 & 4).

**Figure 16:**
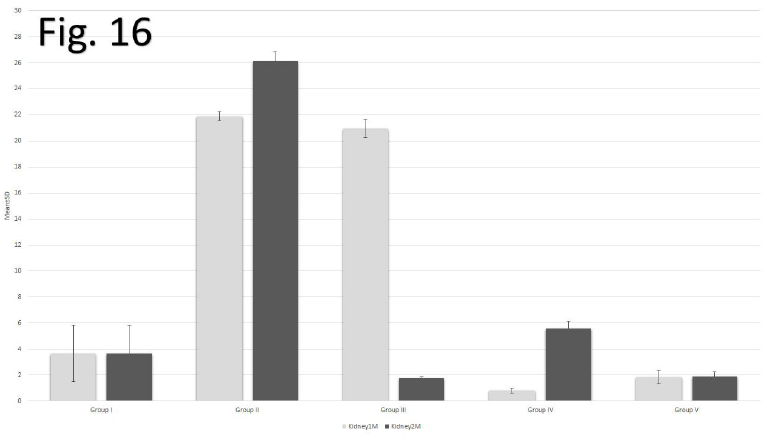
A chart showed the comparison between collagen surface area (Masson Trichrom stain) in renal cortex among studied control & experimental groups in one and two months.

In histopathological examination of Group II (Non-treated Hypertensive rats), myocardium and renal cortex showed marked significant increase in interstitial fibrosis in comparison with that of the same group for one month. (Figs 3, 15 & 16) and (Tables 1 & 2). The anti-VEGF-A antibody stained sections showed significant decrease in the number of cardiac muscle fibers with positive expression of VEGF-A in comparison with that of the control group but it was non-significant in comparison with one month hypertension duration of the same group. While, there was marked significant decrease in the number of cells of renal cortex with positive expression of VEGF-A nearly to the half in comparison with that of the hypertensive rats of the same group for one month. (Figs 11, 17 & 18) and (Tables 3 & 4).

**Figure 3:**
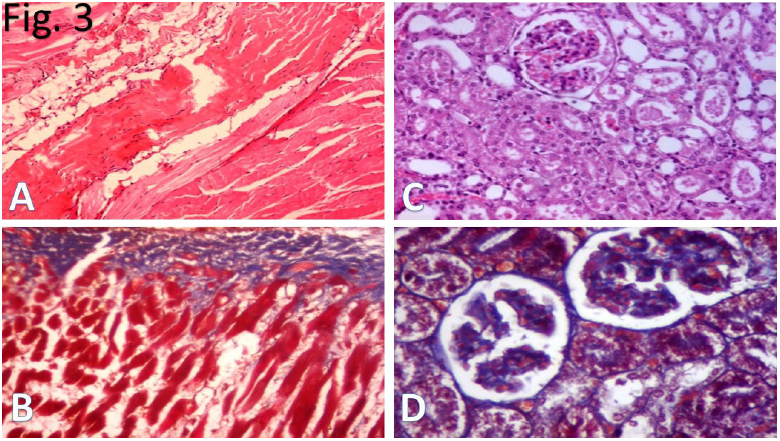
**(A, B, C & D):** Photomicrographs of histological sections of heart (A & B) and kidney (C & D) of Non-treated Hypertensive rats (Group II) for two month. **A & B:** cardiac muscle showed marked increase in intra-myocardial fibrosis (**Hx. & E., X100 & Masson trichrome, X200**). **C & D:** renal cortex showed marked increase in intra-glomerular fibrosis with thickened glomerular and tubular basement membrane (**Hx. & E., X100 & Masson trichrome, X200**).

In Group III (Allopurinol-treated Hypertensive rats), the histopathology of myocardium showed high significant decrease in the surface area of interstitial fibrosis in myocardium and high marked decrease in interstitial fibrosis in the renal cortex in comparison to that of the control, hypertensive non-treated and one month allopurinol-treated hypertensive rats groups. (Figs 5, 15 & 16) and (Tables 1 & 2). This group showed high significant decrease in the maximum number of the myocardium with positive expression of VEGF-A in comparison to groups I & II. The renal cortex still had shown significant decrease in the number of cells with positive expression of VEGF-A in comparison to group I but it showed significant increase in the number of positive expression of VEGF-A in comparison to group II & group III for one month. (Figs 12, 17 & 18) and (Tables 3 & 4).

**Figure 5:**
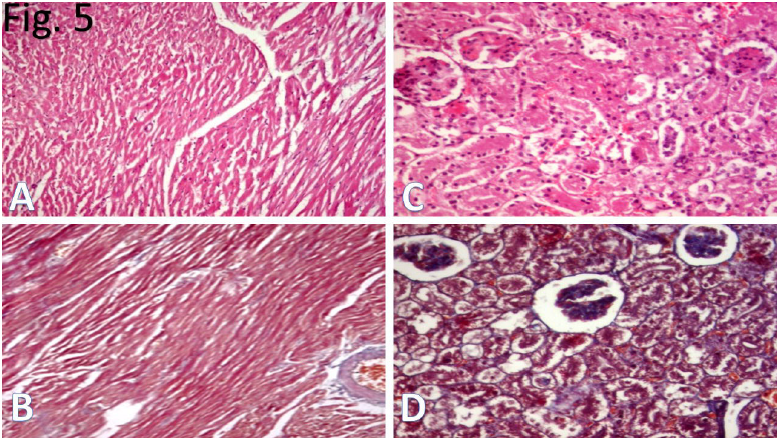
**(A, B, C & D)** Photomicrographs of histological sections of heart (A & B) and kidney (C & D) of Allopurinol-treated Hypertensive rats (Group III) for two month. **A:** cardiac muscle showed disarray of cardiac muscle fibers **(Hx. & E., X100). B:** cardiac muscle showed decreased interstitial fibrosis. **(Masson trichrome, X200)**. **C:** showed renal cortex with ischemic degenerative changes in glomeruli and renal tubules **(Hx. & E., X100) D:** renal cortex showed marked decrease in interstitial fibrosis and thickened glomerular basement membrane **(Masson trichrome, X200)**.

In Captopril-treated Hypertensive rats (Group IV), myocardium showed the most high marked significant reduction in interstitial fibrosis in comparison to groups II & III of the same duration. The renal cortex showed interstitial fibrosis slightly higher than that the control group (group I) (Figs 7, 15 & 16) and (Tables 1 & 2). These results were homogenous with that of the expression of VEGF-A in both myocardium and renal cortex. They showed the most highly significant increase in number of cells with positive expression of VEGF-A in comparison to group II & group III (Figs 13, 17 & 18) and (Tables 3 & 4).

**Figure 7:**
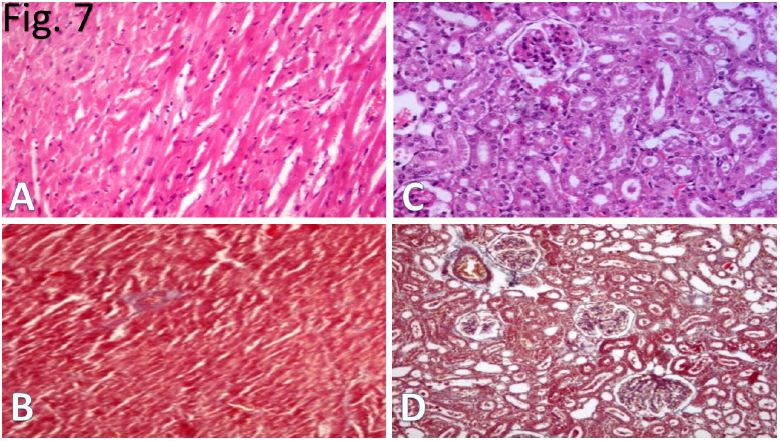
**(A, B, C & D)** Photomicrographs of histological sections of heart (A & B) and kidney (C & D) of Captopril-treated Hypertensive rats (Group IV) for two month. **A:** cardiac muscle became more regular in arrangement **(Hx. & E., X100). B:** cardiac muscle showed diminished interstitial fibrosis **(Masson trichrome, X200). C:** renal cortex showed atrophy of some glomeruli **(Hx. & E., X100) D:** with mild increase in interstitial fibrosis and thickened basement membrane **(Masson trichrome, X100).**

In Group V (Allopurinol-Captopril-treated Hypertensive rats), we studied the combined effects of both allopurinol and captopril. The histopathological examination of both myocardium and renal cortex showed nearly normal architecture as that of the control. They showed the high marked significant reduction in interstitial fibrosis in comparison to that of group IV. The renal cortex showed significant reduction in the surface area of interstitial fibrosis to nearly the half of that of the control group (group I) and the fifth of that of Group IV (Allopurinol-treated Hypertensive rats) (Figs 9, 15 & 16) and (Tables 1 & 2). There were significant improvement in the process of angiogenesis better than that of groups III & IV (Figs 14, 17 & 18) and (Tables 3 & 4).

**Figure 9:**
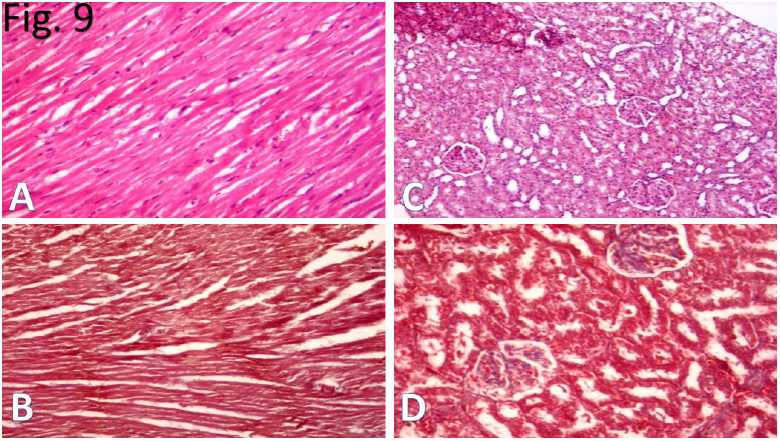
**(A, B, C & D)** Photomicrographs of histological sections of heart (A & B) and kidney (C & D) of Allopurinol-Captopril-treated Hypertensive rats (Group V) for two month. **A & B:** cardiac muscle fibers became more regular in arrangement approximating that of the control group with marked decrease in fibrosis. **C & D:** renal cortex showed normal structure of and with normal glomeruli with minimal fibrosis. **(Hx. & E., X200 & Masson trichrome, X200)**

## 4. Discussion

Hypertension is risk factor for development of congestive heart failure. The pathogenesis of myocardial changes in hypertension includes structural remodeling and fibrosis. Activation of renin-angiotensin system is a key contributing factor of hypertension, and thus interventions that antagonize this systems allows the regression of hypertrophy and heart failure.^29^

Renal oxidative stress appears in hypertensive kidney disease. Fibrosis of the glomerulus and the tubule-interstitium occurs in animals with hypertension.^30^

The early histopathological examination of Group II (Non-treated Hypertensive rats), myocardium and renal cortex showed significant increase in interstitial fibrosis indicated glomerulosclerosis. The anti-VEGF-A antibody stained sections, non-significant increase in the number of cardiac muscle fibers with positive expression of VEGF-A with high significant decrease in the number of positive cells of renal cortex. Late in group II, the myocardium and renal cortex showed marked significant increase in interstitial fibrosis indicating that the hypertension induced duration dependent interstitial fibrosis in both myocardium and renal cortex. The anti-VEGF-A antibody stained sections showed significant decrease in the number of cardiac muscle fibers with positive expression of VEGF-A. These results indicated that the expression of VEGF-A in myocardium was related to the direct effect of hypertension more than its duration. While, there was marked significant decrease in the number of cells of renal cortex with positive expression of VEGF-A nearly to the half in comparison with that of the hypertensive rats of the same group for one month. These results indicated that the expression of VEGF-A in renal cortex was related to both the direct effect and the duration of hypertension. From this group, we concluded that the untreated cases with hypertension showed increased interstitial fibrosis and decreased the expression of VEGF-A in myocardium and renal cortex either through its direct effect or its long course duration with poor angiogenesis.

The effects of ROS on the neonatal rat cardiac fibroblasts can cause both a decrease in fibrillar collagen synthesis and an increase in matrix metalloproteinases (MMP) activity. These results suggested that MMP play an important role in the pathophysiology of myocardial extracellular matrix remodeling during various physiological and pathological conditions.^31^

Evidence proposes that ROS play a key role in the pathophysiological processes of hypertensive renal diseases. Regarding glomerular alterations, ROS mediates lipoprotein glomerulopathy and other inflammatory glomerular lesions.^32^

Other research confirmed marked increased in type I collagen gene expression in glomeruli and interstitial space in AngII induced hypertensive rats. Enhanced renal collagen mRNA is associated with myofibroblasts and accumulated collagen. These observations indicate that collagen synthesis is upregulated in the kidney after oxidative stress induced by Ang-II infusion.^33^

Renal changes associated with hypertension were explained by several studies. Mechanical forces associated with hypertension increased ROS production. ROS-induced vasoconstriction results from increased intracellular calcium concentration, thereby contributing to the pathogenesis of hypertension. ^34^ Other studies explained renal changes by the production of vascular superoxide (NOX) which is derived primarily from NADPH oxidase when stimulated by angiotensin II which is already elevated in hypertension. The activation and production NOX is an important molecular mechanism triggering oxidative injury of podocytes. This may represent an early event initiating glomerulosclerosis and hypertrophy of renal tubular cells. ^35^

Another explanation is homocysteine, is a molecule may play an important role in the pathogenesis of essential hypertension. ^36^ Elevated homocysteinemia diminishes the vasodilation, increases oxidative stress, stimulates the proliferation of vascular smooth muscle cells (VSMC), and alters the elastic properties of the vascular wall.^37^ Elevated homocysteine levels could lead to oxidant injury of the endothelium.^38^

Hypertension-induced ROS stimulate the induction of VEGF expression in various cell types, such as endothelial cells and smooth muscle cells, whereas VEGF induces endothelial cell migration and proliferation through an increase of intracellular ROS. ^39^ A number of studies have demonstrated this positive relationship between ROS and angiogenesis. Hydrogen peroxide induces VEGF expression in vascular smooth muscle cells, as well as endothelial cells, and thereby promotes angiogenic responses.^40^

We compared between the individual effects of allopurinol in Group III and Captopril in Group IV in order to evaluate the effect of combined administration of allopurinol and captopril in group V.

In Allopurinol-treated Hypertensive rats for one month (Group III), the histopathology of myocardium and renal cortex showed high significant increase in interstitial fibrosis. Non-significant number of the cardiac muscle fibers showed positive expression of VEGF-A, renal cortex still had shown significant decrease in the number of cells with positive expression of VEGF-A. After two month, it showed high significant decrease in the surface area of interstitial fibrosis in myocardium and renal cortex. These results confirmed that the long treatment course of hypertension with allopurinol antagonized the fibrotic effects of hypertension in myocardium and renal cortex. This group showed high significant decrease in the maximum number of the myocardium with positive expression of VEGF-A. The renal cortex still had shown significant decrease in the number of cells with positive expression of VEGF-A. The study of VEGF-A in this group concluded that the allopurinol had a dual effects as it decreased its expression in the myocardium and increased its expression in the renal cortex.

Very High dose of Allopurinol (up to 50 mg/kg/day) acted directly as a scavenger for the free radicals with antioxidant properties as demonstrated in vitro hearts. ^41^ On the other hand, other researches evidenced that lower doses of allopurinol (sufficient to block XO activity) failed to show antioxidant protection but higher doses did. ^42^

Another study that used both very high dose of allopurinol (100 mg/day for two weeks) and myocardial infarction inducer isoproterenol (ISO) (at a dose of 50mg/kg twice a week for two weeks) reveals that allopurinol exerts significant cardio-protective effect against ISO induced myocardial infarction in aged rats. This protective effect could be associated with the enhancement of antioxidant defense system and attenuation of inflammatory cells infiltration in the myocardium.^43^

The mechanism of improvement in endothelial function with high dose of allopurinol lies in its ability to profound reduction in vascular oxidative stress and not in urate reduction.^44^

The results of other research obtained from non-hyperuricemic obstructed kidney model concluded that increase in Xanthine oxidase (XO) activity itself may play an important role in the progression of tissue fibrosis. An example for Xanthine oxidase (XO) inhibitor (febuxostat), may be a therapeutic tool for treating progressive interstitial fibrosis.^45^

Allopurinol exhibited dual effects (angio-preventive and angio-promoting effects) in VEGF gene expression on the experimental inflamed fibro-vascular tissue induced in mice by synthetic subcutaneous implant. The contrasting findings were dependent on the phase in which the treatment was initiated. An angio-preventive effect (same as control / decrease in VEGF level) was observed when treated with allopurinol during acute inflammation, whereas angio-promoting effect (increase in VEGF level) was seen treated with allopurinol during the chronic process (that was initiated 8 days post-implantation).^46^

In comparison, early Captopril-treated Hypertensive rats (Group IV), myocardium and renal cortex still showed marked significant reduction in interstitial fibrosis. As in groups I, II & III, the number cardiac muscle fibers with positive expression of VEGF-A was still non-significant. The renal cortex showed significant decrease in the number of cells with positive expression of VEGF-A. Late in this group, myocardium showed the most high marked significant reduction in interstitial fibrosis. This indicated that the captopril had more powerful anti-fibrotic effect than that of allopurinol alone through its direct effects and long course treatment. The renal cortex showed interstitial fibrosis slightly higher than that the control group. These results were homogenous with that of the expression of VEGF-A in both myocardium and renal cortex. This group especially this duration (two month) showed the most highly significant increase in number of cells with positive expression of VEGF-A. This means that the captopril alone improved the angiogenesis of myocardium and renal cortex even with the long course of hypertension. This ment that the captopril prevented the progression of the renal impairment. This indicated that the captopril had protective renovascular effects.

The oxidative stress in patients with essential hypertension was improved by chronic administration of therapeutic doses of the sulfhydryl ACE inhibitor but not with the nonsulfhydryl ACE inhibitor. By these data, authors concluded the protective effect of the sulfhydryl ACE inhibition in retarding vascular dysfunction and atherogenesis that often develops rapidly in hypertensive patients.^47^

ACE inhibitors as antioxidant strategy had been prooved through ameliorate vasoconstriction, increase the bioactivity of NO, and can inhibit vascular superoxide production at its source. This is why ACE inhibitors may represent a “magic bullet” against vascular oxidative stress.^48^

The ACE inhibitors had dual effects in the process of angiogenesis. They improved neovascularization in the diabetic ischemic leg through activation of bradykinin signaling, whereas it reduced vessel growth in the diabetic retina through inhibition of overacting Ang II pathway.^49^

Also, it was observed that the addition of ACE inhibitors to endothelial progenitor cells therapy induced the neovascularization and it reduced the number of apoptotic cardiomyocytes.^50^

In Group V Combined Allopurinol-Captopril-treated Hypertensive rats for one month, myocardium and renal cortex showed significant reduction in interstitial fibrosis. The immunohistochemical stained sections, there was non-significant increase in the number of cardiac muscle fibers with positive expression of VEGF-A. The renal cortex still had showed significant decrease in the number of cells with positive expression of VEGF-A. After two month, both myocardium and renal cortex showed nearly normal architecture as that of the control. These results was confirmed by Masson trichrome stained sections in which they showed the high marked significant reduction in interstitial fibrosis in comparison to that of group IV. These indicated that the combination between captopril and allopurinol had the same powerful anti-fibrotic effect as that of captopril alone through their direct effects and long course treatment. But, the renal cortex showed significant reduction in the surface area of interstitial fibrosis to nearly the half of that of the control group (group I) and the fifth of that of Group IV (Allopurinol-treated Hypertensive rats). This is why it is better to subscribe the combination between the allopurinol and captopril than that to subscribe one of them alone because this combination was nephron-protective. Moreover, immunohistochemical stained sections added more evidence for the benefits of this combination. In which, there was significant improvement in the process of angiogenesis.

Animal studies proved that the captopril and allopurinol in combination preserved normal blood pressure and insulin sensitivity and prevented hypertriglyceridemia, hyperuricemia, and hypercholesterolemia. The exploration of potential mechanisms that resulted in a superior effect of combined therapy will be evaluated in future studies.^51^

Researches on human concluded that the long term maintenance of captopril and allopurinol can lead to severe adverse effects. There have been a small number of cases of Stevens-Johnson syndrome. The combination of ACE inhibitors and allopurinol can also increase the risk of blood dyscrasias (eg, leucopenia, neutropenia), resulting in serious infection, and patients with renal impairment are at greater risk. However, these adverse effects are rare and unpredictable, and do not preclude the use of both medicines.^52^

## 5. Summary and Conclusion

Experimentally, long term therapy with the combination between allopurinol and captopril is of high benefit for both myocardium and renal cortex in case of chronic hypertension. As the combination decreases the fibrotic changes associated with hypertension and enhances the process of angiogenesis.

